# Assessment of the level of pollution based on soil and *Tilia × europaea* leaves

**DOI:** 10.1101/2024.04.17.589890

**Authors:** Edina Simon, Haziq Bin Ismail, Bianka Sipos, Vanda Éva Abriha-Molnár, Dávid Abriha, Dávid Tőzsér, Zsófi Sajtos, Rafael Boluda, Luis Roca-Pérez

## Abstract

Anthropic activities related to urbanization release pollutants, including potentially toxic elements, into the atmosphere and these are eventually deposited in soils, water, infrastructures, vegetation. Urban soil and plant leaves are widely used as ecological indicators to assess the effect of urbanization. This study aimed to evaluate the impact of urbanization based on the elemental concentration of soil and leaves of *Tilia × europaea* from urban, suburban, and rural areas along an urbanization gradient in Debrecen, Hungary. Using the ICP-OES technique, we measured the concentration of Ba, Cd, Co, Cr, Cu, Ni, Pb, Sr, and Zn and based on the measured concentration, bioaccumulation factor (BAF) was calculated. The highest concentration of all elements was found in soil samples from urban areas, with an increasing tendency along the urbanization gradient. A significant difference was found along the urbanization gradient only for Cr based on the plant leaves. *T. × europaea* showed bioaccumulation capacity for Cr and Sr. Our results suggest that urbanization has a remarkable effect on the elemental concentration of soil, which is a perfect ecological indicator. At the same time, we concluded that the *T. × europaea* was not sensitive enough to indicate the effect of urbanization.

## Introduction

Soil and air pollution by metals is one of the most critical environmental problems worldwide (Hu et al. 2013). Anthropic activities cause contamination in soils by certain elements such as Sr, As, Cd, Cr, Cu, Ni, Pb and Zn (Roca-Perez et al. 2010); among anthropogenic activities, urbanization is one of the primary sources of polluted soil with heavy metals (Mirales et al. 2012). Urbanization is progressing rapidly between 2000 and 2030, and the urban population is anticipated to grow an average of 2.3% per year in the developing world (Chen 2007). The effects of urbanization on environmental pollution are different in countries and regions with various degrees of development (Son and Hu 2018). Earlier studies reported that urbanization directly and indirectly affects soil as an essential resource; accumulation of heavy metals can modify the quality of soil and agricultural products, and thus, they have adverse effects on humans, animals, and the ecosystem health (Nagajyoti et al. 2010, Ihenetu et al. 2024)

Trees play an essential role in urban vegetated areas (Ma et al. 2022, Wang and Zhang 2022). Plant leaves can accumulate air pollutants in high concentrations; air pollutant particles can also be deposited in the leaves of plants due to their stomatal openings and waxy cuticles (Bibi et al. 2023, Molnár et al. 2020, Simon et al. 2021). The genus Tilia is generally used as biomonitors of trace elements to assess the effect of urbanization and industrialization on terrestrial environments (Zeb et al. 2022, Andrianjara et al. 2024).

The bioaccumulation factor determines the capacity of plants to transfer and accumulate heavy metals or other compounds from soils to the aerial part of the plant (Hosseini et al. 2022; Korzeniowska & Stanislawska-Glubiak 2019, Roca-Perez et al. 2023). This index has been applied to soil-plant system in mining areas (Roca-Perez et al. 2023) near road networks (Hosseini et al. 2022), rural and urban areas (Pietrelli et al. 2023); in urban areas with species of the genus *Tilia* (Mitrović et al. 2021; Mitrovi et al. 2023).

Therefore, *Tilia × europaea* was chosen for our study, a typically ornamental tree in European countries (Ţenche-Constantinescu et al. 2015). Our study aimed to investigate elemental concentration in soil and *Tilia × europaea* plant leaves along an urbanization gradient in Debrecen. Hungary. Our hypotheses are as follows: (i) there are significant differences in pollutant levels along the urbanization gradient (urban, suburban, and rural areas), (ii) the highest metal concentration in soil and leaves is in the urban area, while the lowest concentration is in the rural area, (iii) significant correlation is found between the soil and leaves metal concentrations.

## Material and methods

### Study area

Debrecen is the second largest city in Hungary, with a population of approximately 200.000 as of 2019. The area allocated for the city is 461 km^2^, which results in a population density of 436/km^2^. Along an urbanization gradient in Debrecen city, three different studied areas were chosen (Fig.1). The rural area was situated near Nagyerdő, a protected area for nature conservation. The suburban area is intermediate between the rural and urban areas regarding urban density. It is located adjacent to a road with apartments on one side, and the side is a forest. Lastly, the urban study area was at the center of the urban community to maximize the observable effect of urban pollution, and it was surrounded by housing and offices at the center of Debrecen. Soil samples (N = 15) were collected along an urbanization gradient representing rural, suburban, and urban areas. At each sampling area, five soil samples were collected. All samples were collected from 0–20 cm depth with a small hand spade. Samples were placed into poly bags and kept at -21 °C in the refrigerator.

**Fig 1.**
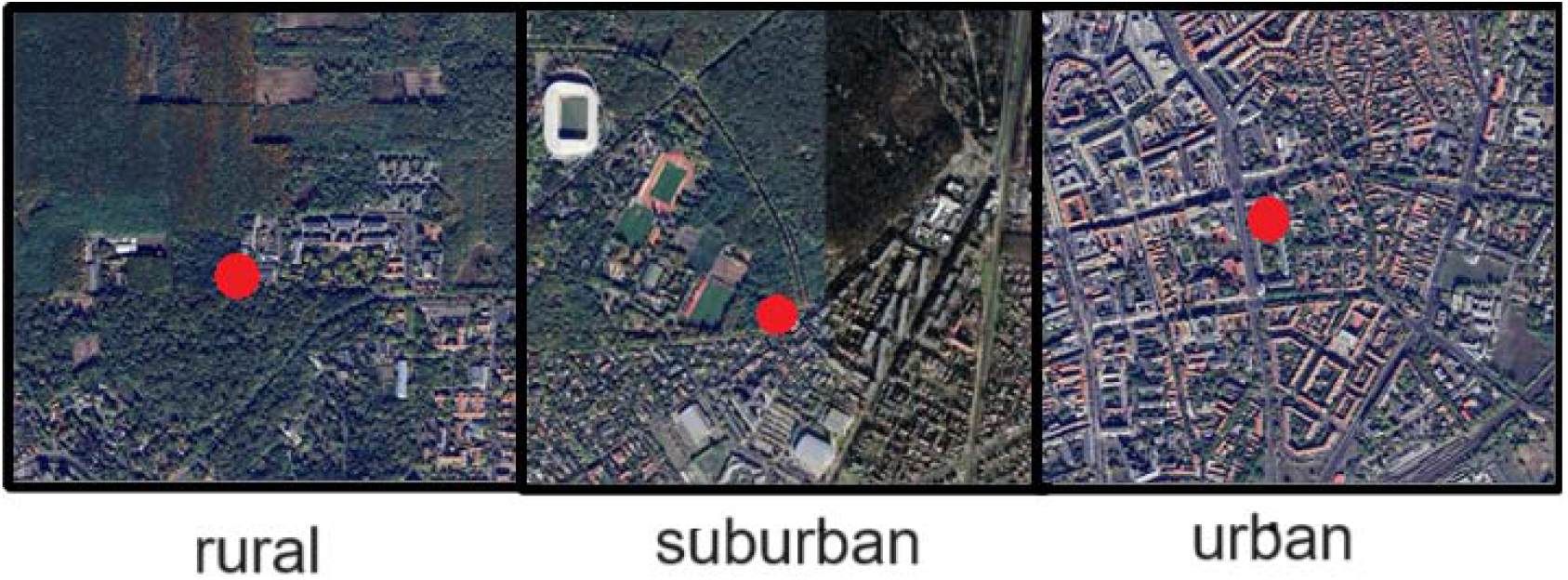
Studied areas for Debrecen along an urbanization gradient. Notations: The red circle indicates the study areas

During the plant sampling, 10-15 leaves were collected from 3 trees of the *Tilia × europaea* in the same location. Additionally, soil samples were collected within a 1-meter radius of the tree base to calculate the bioaccumulation factor. Trees showing any form of disease should be avoided, as this may alter the investigation results.

#### Sample preparation

The laboratory process started with a surface wash to remove any surface pollutants from the leaf samples. After washing, the leaf samples were dried in an oven set at 70 °C for 90 minutes. Soil samples were dried at room temperature, and after that stones, plant roots, and residues were removed with plastic tweezers. Samples were sieved in a 2-mm plastic sieve. Then, the sample process was the same for soil and leaf samples. The samples process was published by Simon et al. (2014, 2016, 2021), Bibi et al. (2023a, b), and Molnár et al. (2020) (Fig. 2). For the elemental analysis, ICP-OES was used. Peach leaves (1547) and soil (SQC001) CRM were used, and the recoveries were within 10% of the certified values for the elements (Simon et al. 2013, 2014, 2016, 2021). The concentration of Ba, Cd, Co, Cr, Cu, Ni, Pb, Sr, and Zn was analyzed during the analysis.

**Fig 2.**
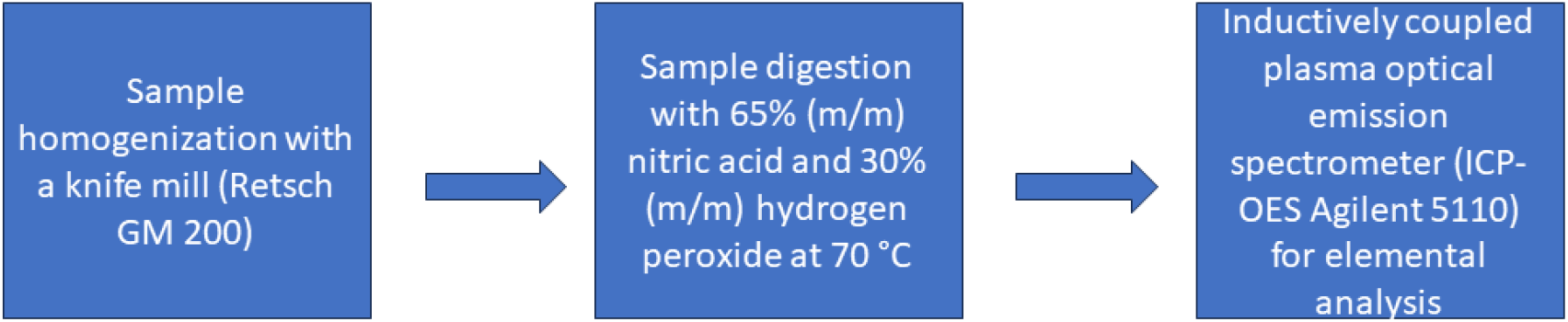
Sample preparations and elemental analysis for soil and leaves

### Bioaccumulation factor (BAF)

To study the accumulation of elements in leaves of *T. × europaea*, bioaccumulation factor (BAF) was used. BAF is the ratio of elemental concentration in leaves (C_plant_) and elemental concentration in soil (C_soil_) (Ivanciuc et al. 2006, van Gestel et al. 2011):

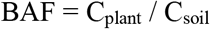

### Statistical analyses

Statistical analyses were conducted with SPSS Statistics 20 (IBM) statistical software. The normality of the distribution was tested using the Shapiro–Wilk test. The homogeneity of variances was tested using Levene’s test. The differences among samples were tested with the analysis of variance (ANOVA) for each variable. Tukey’s test was performed for pairwise comparison between the groups. Canonical discriminant analysis (CDA) was used to reduce dimensions and to identify those variables that most efficiently discriminated the study area as the dependent variable (De Sá 2007). The correlation between the elemental concentration of soil and leaves was analyzed with the Pearson correlation coefficient (Zar 1996).

## Results

There was a clear separation among the studied areas based on the elemental concentration of soil according to the CDA (Fig. 3). The canonical variables were significant (*p* < 0.001) in the first and second discriminant functions. The percentage of variance was 87.0 in the first, while 13.0 was in the second discriminant function. There was significantly higher elemental soil concentration in the urban area than in the rural area (Table 1). The order of contents of potentially toxics elements was: Zn>Ni>Ba>Pb=Sr>Cu>Cr>Co>Cd in rural soil, Ba>Zn>Ni>Sr>Pb>Cu>Cr>Co>Cd in suburban soils, and Ba>Zn>Ni>Sr>Cr>Pb >Cu>Co>Cd in urban.

**Table 1.** Summary statistics of elemental concentration of soil (mg kg^-1^, mean ± SE).

| Elements | Urbanization gradient |  |  | F | p |
| --- | --- | --- | --- | --- | --- |
|  | rural | suburban | urban |  |  |
| Ba | 14.2 ± 1.0 | 67.7 ± 16.2 | 97.1 ± 11.0 | 13.825 | 0.001 |
| Cd | 0.3 ± 0.1 | 0.8 ± 0.1 | 1.8 ± 0.2 | 55.623 | <0.001 |
| Co | 1.3 ± 0.1 | 2.6 ± 0.1 | 6.4 ± 0.4 | 96.986 | <0.001 |
| Cr | 3.1 ± 0.3 | 7.3 ± 0.7 | 28.0 ± 6.3 | 13.081 | 0.001 |
| Cu | 3.7 ± 0.3 | 13.8 ± 2.5 | 22.3 ± 2.7 | 19.031 | <0.001 |
| Ni | 18.9 ± 4.5 | 28.6 ± 1.5 | 39.8 ± 4.2 | 8.229 | 0.006 |
| Pb | 5.5 ± 2.3 | 20.9 ± 5.5 | 24.5 ± 3.9 | 5.971 | 0.016 |
| Sr | 5.5 ± 0.7 | 25.2 ± 6.8 | 32.6 ± 4.3 | 8.975 | 0.004 |
| Zn | 19.1 ± 1.2 | 61.5 ± 8.1 | 83.9 ± 13.7 | 12.735 | 0.001 |

**Fig 3.**
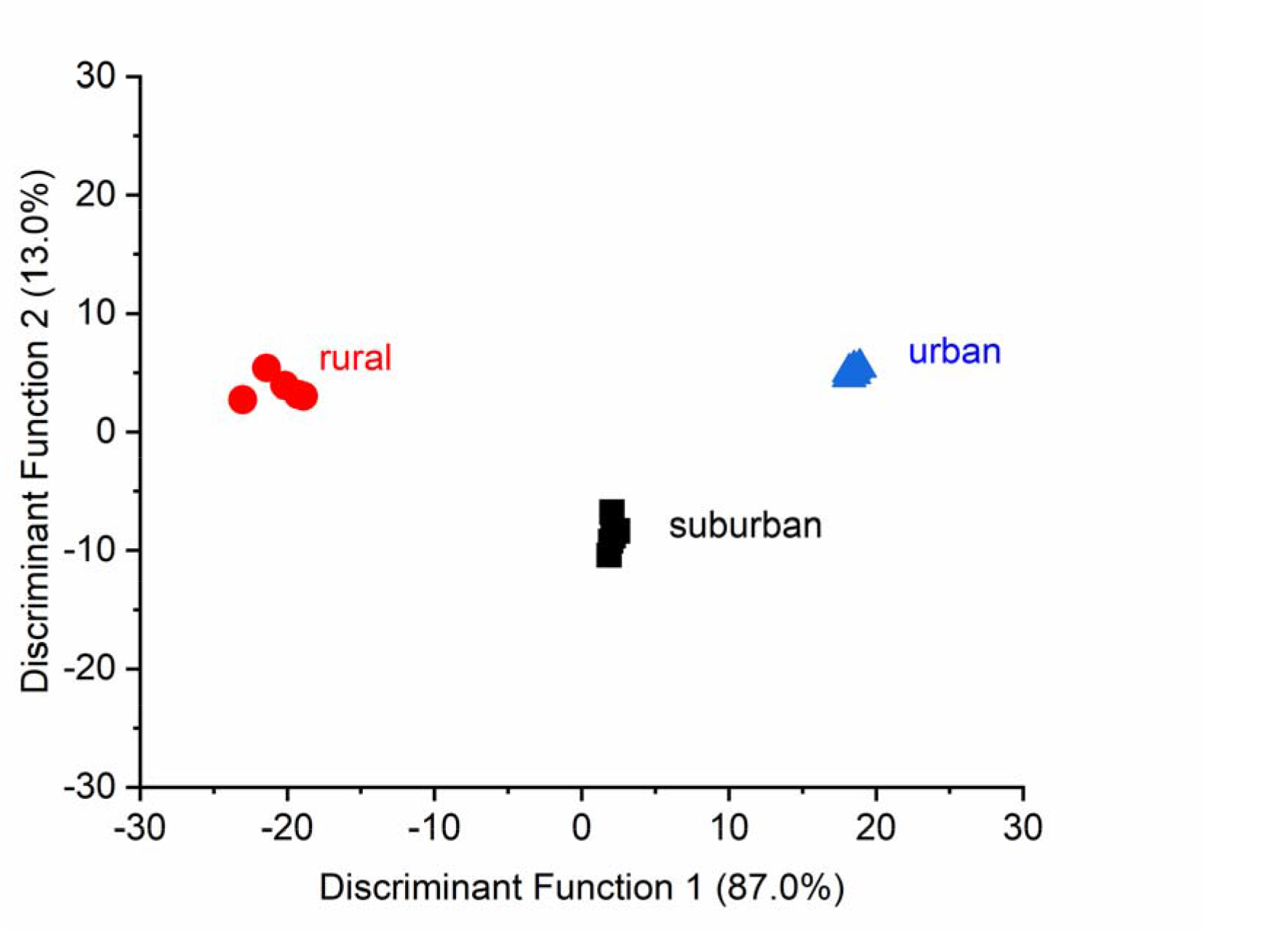
Scatter plot of canonical discriminant analysis (CDA) based on the elemental concentration of soil among the studied areas

Similar to the soil, separation was found among the studied areas based on the elemental concentration of *Tilia × europaea* according to the CDA (Fig. 4). The canonical variables were not significant (*p* > 0.05) in the first and second discriminant functions. The percentage of variance was 99.2 in the first, while 0.8 was in the second discriminant function. In rural areas, higher concentrations of Ba, Cu, Ni and Zn were observed in the leaves of *T*.*× europaea*; however, only Cr in rural and suburban areas showed significantly higher values than in urban areas (Table 2). The Cd, Co, Cr, and Pb concentrations were below the detection limit.

**Table 2.** Summary statistics of elemental concentration of *Tilia × europaea* leaves (mg kg^-1^, mean ± SE).

| Elements | Urbanization gradient |  |  | F | p |
| --- | --- | --- | --- | --- | --- |
|  | rural | suburban | urban |  |  |
| Ba | 12.9 ± 4.2 | 8.1 ± 0.6 | 5.6 ± 2.5 | 1.663 | 0.266 |
| Cr | 0.3 ± 0.1 | 0.3 ± 0.1 | 0.2 ± 0.1 | 7.426 | 0.024 |
| Cu | 7.7 ± 0.9 | 6.2 ± 0.6 | 4.9 ± 1.1 | 2.709 | 0.145 |
| Ni | 1.5 ± 0.5 | 0.6 ± 0.1 | 0.5 ± 0.1 | 3.844 | 0.084 |
| Sr | 28.8 ± 4.6 | 41.9 ± 1.7 | 32.1 ± 5.8 | 2.456 | 0.166 |
| Zn | 18.5 ± 1.4 | 14.4 ± 0.4 | 17.5 ± 0.9 | 4.85 | 0.056 |

**Fig 4.**
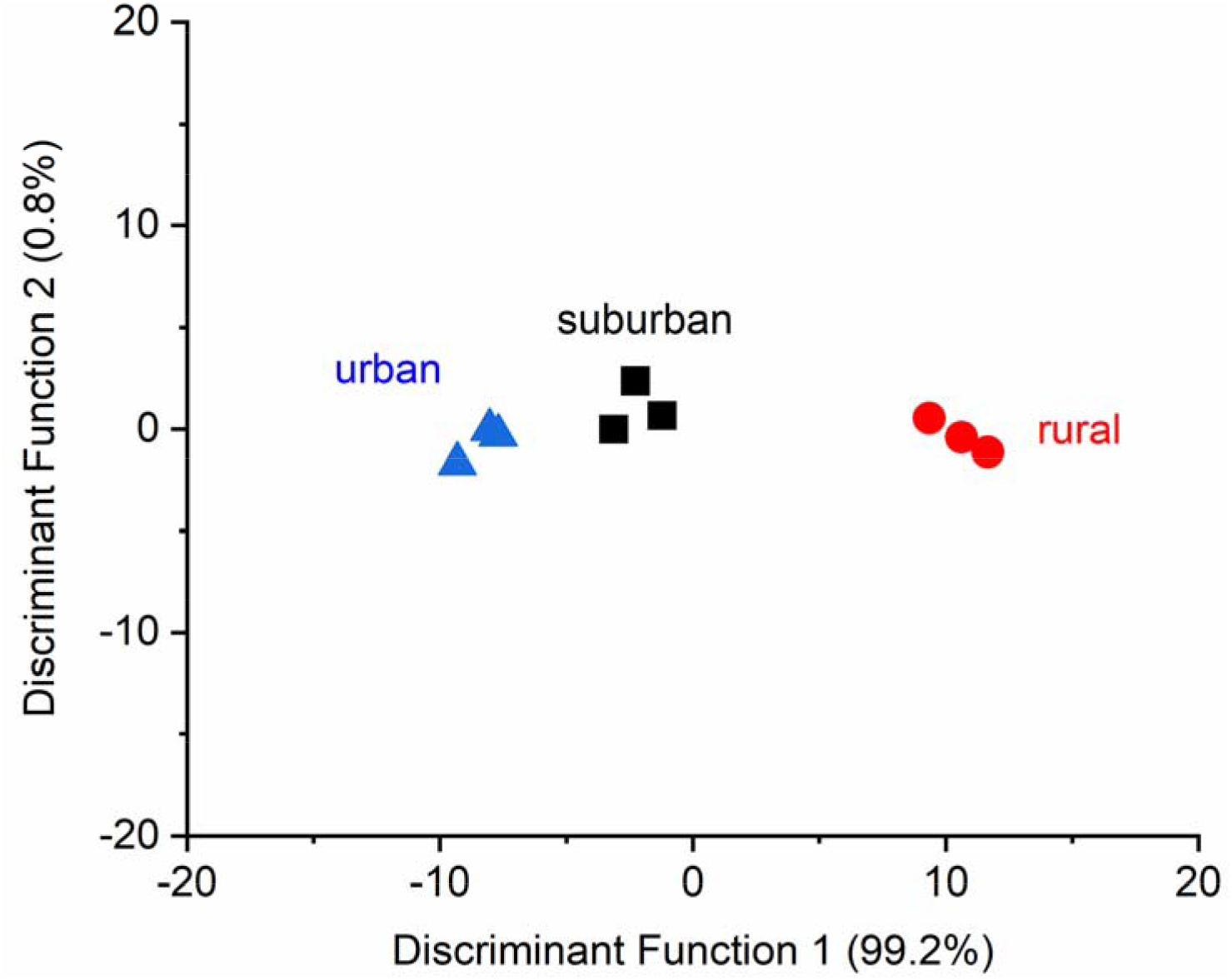
Scatter plot of canonical discriminant analysis (CDA) based on the elemental concentration of *Tilia × europaea* leaves among the studied areas

In all studied areas, we did not find any significant correlation between the elemental concentration of soil and leaves (rural: r = 0.429, *p* = 0.397; suburban: r = 0.486, *p* = 0.329; urban: r = 0.371, *p* = 0.468). The BAF values were higher than 1 for Cu in the rural and for Sr in all studied areas (Fig. 5 and 6). The BAF values of Cu and Sr indicated that trees could accumulate these elements from soil indirectly or directly.

**Fig 5.**
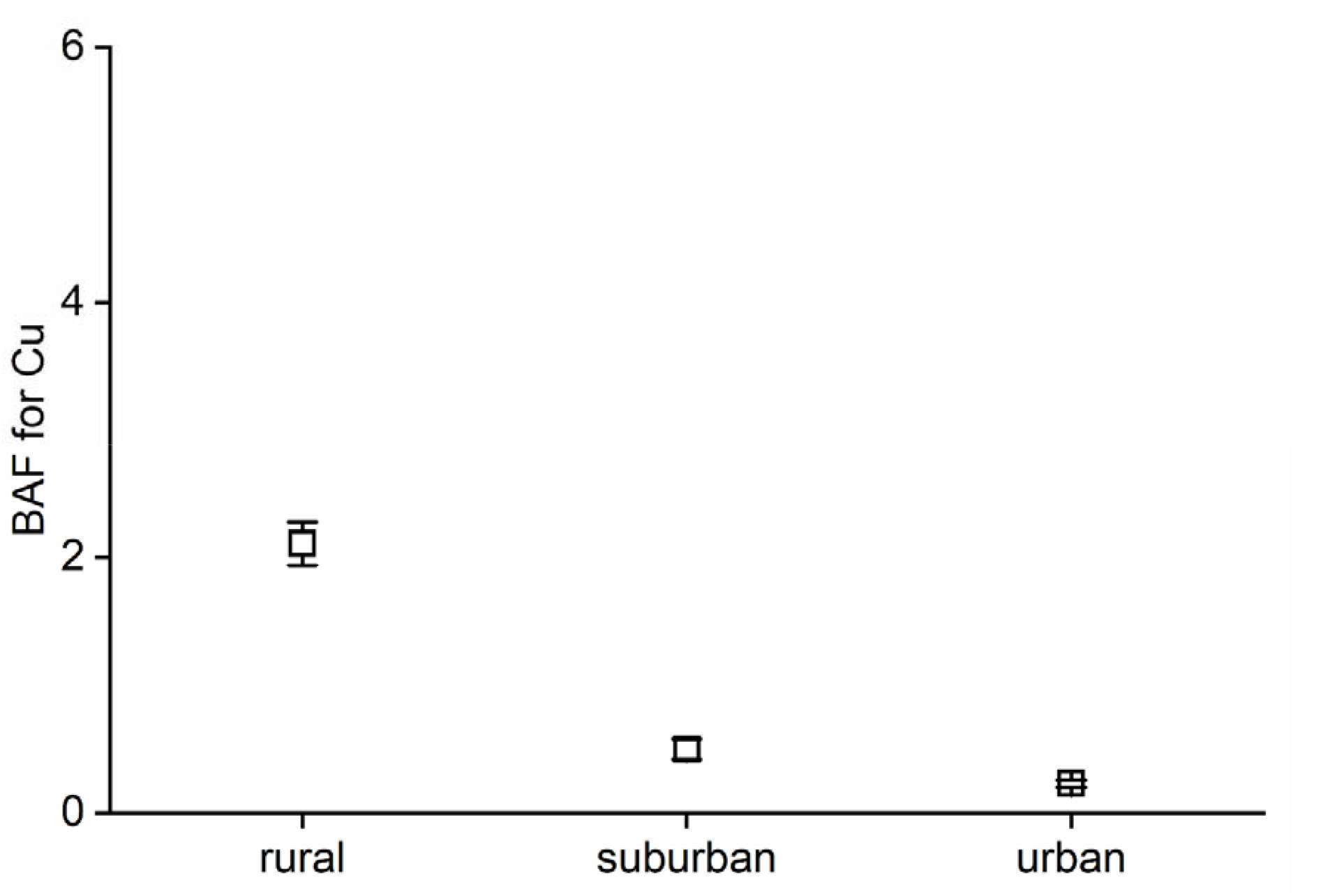
Value of bioaccumulation factor (BAF) for Cu (mean ± SE) along the urbanization gradient

**Fig 6.**
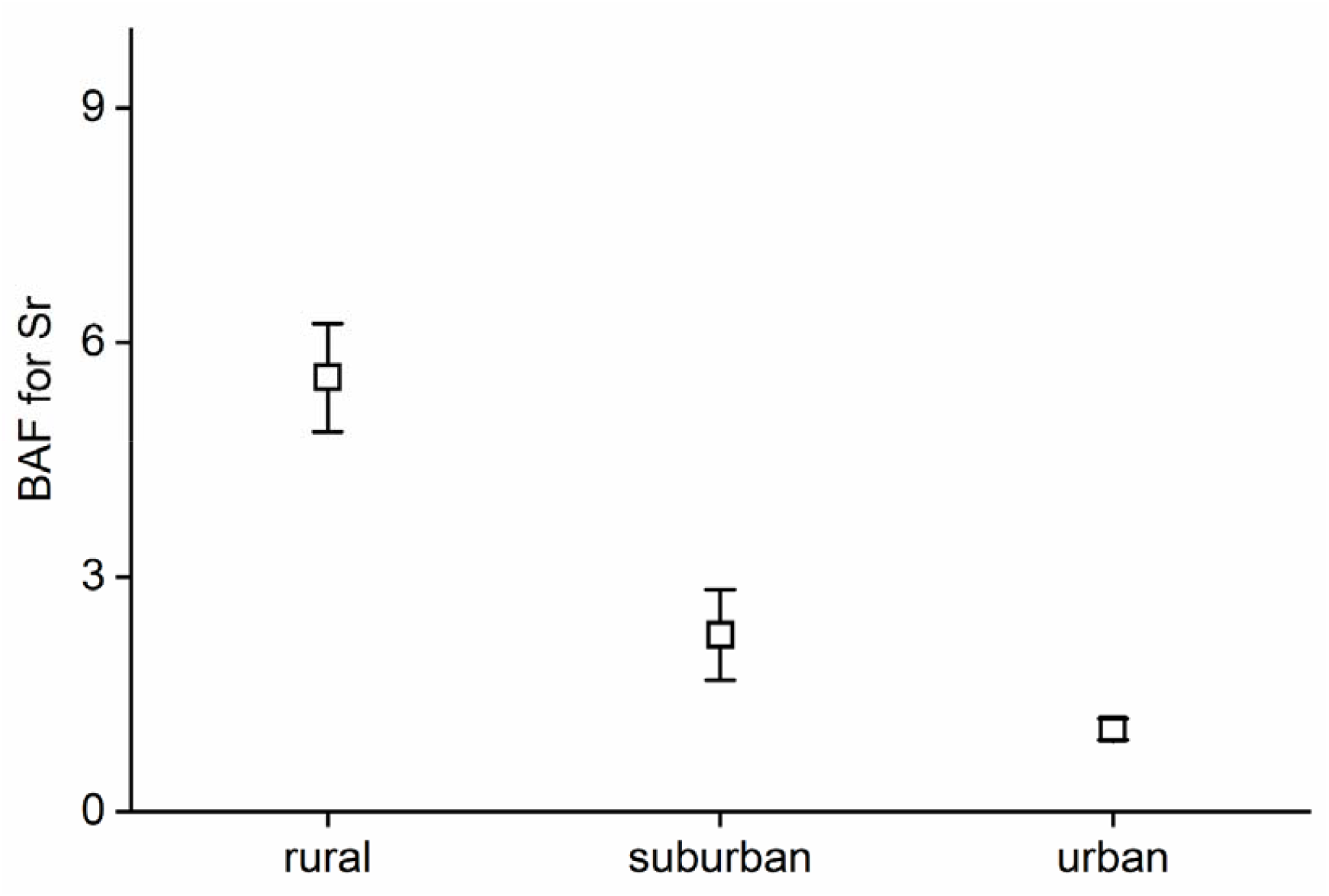
Value of bioaccumulation factor (BAF) for Sr (mean ± SE) along the urbanization gradient

## Discussion

As hypothesized, we found differences based on the elemental concentration of soil along the urbanization gradient; significantly higher concentrations were found in urban areas than in rural ones. Thus, it was indicated that urbanization had a remarkable effect on the metal concentration of soil. Contrary to our hypothesis, the elemental concentration of *T. × europaea* did not differ significantly along the urbanization gradient, only the concentration of Cr, which is a typically anthropogenic element. At the same time, our results also indicated that *T. × europaea* was not sensitive enough to reveal the effect of urbanization.

Similar to our results, Hu et al. (2013) also demonstrated the effect of urbanization on the soil elemental concentration change, and they found the highest pollution level and potential ecological risk of surface soil polluted with metals in urban soil. Urbanization may cause an increasing risk of soil pollutants through waste disposal and acid deposition derived from urban air pollution (Chen 2007). Tešic et al. (2022) also found significantly higher Zn, Cu, Cd, Cr, and Ni in the soil of urban areas in Belgrade than in the rural. Simon et al. (2016) also reported that soil was a suitable bioindicator for monitoring the effects of urbanization and anthropogenic activities on terrestrial ecosystems because they found significant differences among areas based on the concentration of Ba, Cu, Mn, Sr, and Zn in soil. Simon et al. (2013) also confirmed that the elemental analysis of soil samples is one of the best ways to study the effects of urbanization. Bibi et al. (2023) also showed higher concentrations of Cu, Pb, Sr, and Zn were in the urban and suburban areas than in the rural areas in Wien, Austria. In summary, as indicated by Wang et al (2024), urbanization intensifies the impact on potential toxic elements in soil. In relation to the higher or lower content of elements in the soil, the elements with the highest concentration in urban area were Ba, Zn and Ni, these results are similar to those performed in urban soils of Vienna (Austria) where the concentrations of Ba and Zn were also among the highest of the elements analyzed (Simon et al. 2013).

Molnár et al. (2020) demonstrated that *T. × europaea* is sensitive to air pollutants based on the air pollution tolerance index (APTI). Serbula et al. (2013) showed that pine needles responded more intensively to pollution than *Tilia*. Serbula et al. (2013) also indicated, based on the bioaccumulation and translocation index, that the high concentrations of Cu and Pb in aerial parts of pine and linden are mainly a consequence of airborne pollution, which explains that there was no correlation between the elemental concentration of soil and leaves in our study. Similarly to our finding, Ţenche-Constantinescu et al. (2015) also demonstrated that *Tilia* sp. was very resistant to biotic and abiotic stress because they found a correlation only between traffic intensity and the concentration of Pb in leaves. The study of *Tilia* leaf showed a different capacity for accumulating potentially toxic elements in various locations by Mitrović et al. (2021) in urban areas with varying pollution levels. Jouraeva et al. (2002) studied the accumulation potential of PAHs and metals on the leaves of *Tilia×euchlora* and *Pyrus calleryana*, and they reported that leaves of *T. ×euchlora* accumulated higher concentrations of air pollutants than *P. calleryana*. Aničić et al. (2011) studied the metal concentration in the *Aesculus hippocastanum* leaves and *Tilia* spp. Similar to our finding, they also demonstrated that *Tilia* spp. was not so sensitive to assessing Pb and Cu atmospheric pollution (Aničić et al. 2011). Urošević et al. (2019) studied the metal concentration of different tree leaves, and they did not recommend *Tilia* sp. as a valuable bioindicator for air pollutants. In the case of plant species, the uptake of the elements may be accompanied by a species-specific pattern of physiological reaction (Drzewiecka et al. 2019).

In this study only Sr, in all areas, and Cu, only in rural areas, presented BAF values higher than 1, which indicates that *Tilia x europaea* has the capacity to accumulate these elements in leaves. However, studies with *Tilia tomentosa* (Mitrovi et al. 2023), *Tilia argentea* (Greksa et al. 2019) and *Tillia sp*. (Alagić et al. 2013), conducted in different areas with anthropic influence, did not obtain BAF values higher than 1 for all the elements analyzed

## Conclusion

Our study demonstrated differences in the elemental concentration of soil along the urbanization gradient. Slight difference was found in elemental concentrations of *T. × europaea* only in the case of Cr along the urbanization gradient. Based on the accumulation rate, *T. × europaea* could accumulate the Cu and Sr from soil directly or indirectly. Thus, we concluded that urbanization has a remarkable effect on the elemental concentration of soil, which is a perfect ecological indicator. At the same time, we concluded that the *T. × europaea* was not sensitive enough to indicate the effect of urbanization.

## Acknowledgements

Research supported by 2021-1.2.4-TÉT-2021-00049 project and the University of Debrecen Program for Scientific Publication. E. Simon thanks funding from the HUN-REN Hungarian Research Network.

## References

Alagić Sč, Šerbula SS, Tošić SB, Pavlović AN, Petrović JV (2013) Bioaccumulation of arsenic and cadmium in birch and lime from the Bor Region. Arch Environ Contam Toxicol 65: 671–682. 10.1007/s00244-013-9948-7

Andrianjara I, Cabassa C, Lata JC, Hansart A, Raynaud X, Renard M, Nold F, Genet G, Planchais S (2024) Characterization of stress indicators in Tilia cordata Mill. as early and long-term stress markers for water availability and trace element contamination in urban environments. Ecol Indic 158:111296. 10.1016/j.ecolind.2023.111296

Aničić M, Spasić T, Tomašević M, Rajšić S, Tasić M (2011) Trace elements accumulation and temporal trends in leaves of urban deciduous trees (Aesculus hippocastanum and Tilia spp.). Ecol Indic 11:824–830. 10.1016/j.ecolind.2010.10.009.

Aničić Urošević M, Jovanović G, Stević N, Deljanin I, Nikolić M, Tomašević M, Samson R (2019) Leaves of common urban tree species (Aesculus hippocastanum, Acer platanoides, Betula pendula and Tilia cordata) as a measure of particle and particle-bound pollution: a 4-year study. Air Qual Atmos Hlth 12:1081–1090. 10.1007/s11869-019-00724-6

Bibi D, Tőzsér D, Sipos B, Molnár VÉ, Tóthmérész B, Simon E (2023b) Complex study of air pollution based on tree species in Vienna. Air Qual Atmos Hlth 17:417–424. 10.1007/s11869-023-01452-8

Bibi D, Tőzsér D, Sipos B, Tóthmérész B, Simon E (2023a) Heavy metal pollution of soil in Vienna, Austria. Water Air Soil Pollut 234:232. 10.1007/s11270-023-06244-5

Chen J (2007) Rapid urbanization in China: A real challenge to soil protection and food security. Catena 69:1–15. 10.1016/j.catena.2006.04.019

De Sá JPM (2007) Applied Statistics Using SPSS. STATISTICA. MATLAB and R. 2nd ed.; Springer: Berlin/Heidelberg. Germany. ISBN 978-3-540-71971-7.

Drzewiecka K, Piechalak A, Goliński P, Gąsecka M, Magdziak Z, Szostek M, Budzyńska S, Niedzielski P, Mlecze M (2019) Differences of Acer platanoides L. and Tilia cordata Mill. Response patterns/survival strategies during cultivation in extremely polluted mining sludge – A pot trial. Chemosphere 229:589–601. 10.1016/j.chemosphere.2019.05.051

Greksa A, Ljevnaić-Mašić B, Grabić J, Benka P, Radonić V, Blagojević B, Sekulić M (2019) Potential of urban trees for mitigating heavy metal pollution in the city of Novi Sad, Serbia. Environ Monit Assess 191: 636. 10.1007/s10661-019-7791-7

Hosseini NS, Sobhanardakani S, Cheraghi M, Lorestani B, Merrikhpour H (2022) Expansive herbaceous species as bio-tools for elements detection in the vicinity of major roads of Hamedan Iran. Int J Environ Sci Tech 19: 1611–1624.

Hu Y, Liu X, Bai J, Shih K, Zeng EY, Cheng H (2013) Assessing heavy metal pollution in the surface soils of a region that had undergone three decades of intense industrialization and urbanization. Environ Sci Poll Res 20:6150–6159. 10.1007/s11356-013-1668-z

Ihenetu SC, Li G, Mo Y, Jacques KJ (2024) Impacts of microplastics and urbanization on soil health: An urgent concern for sustainable development. Green Anal Chem 8:100095. 10.1016/j.greeac.2024.100095

Ivanciuc T, Ivanciuc O, Klein DJ (2006) Modeling the bioconcentration factors andbioaccumulation factors of polychlorinated biphenyls with posetic quantitativesuperstructure/activity relationships (QSSAR). Mol Diver 10:133–145.

Jouraeva VA, Johnson DL, Hassett JP, Nowak DJ (2002) Differences in accumulation of PAHs and metals on the leaves of Tilia×euchlora and Pyrus calleryana. Environ Poll 120:331–338. 10.1016/S0269-7491(02)00121-5

Korzeniowsk J, Stanislawska-Glubiak E (2019) Phytoremediation potential of Phalaris arundinacea, Salix viminalis and Zea mays for nickel-contaminated soils. Int J Environ Sci Tech 16: 1999–2008.

Ma Z, Zhang P, Hu N, Wang G, Dong Y, Guo Y, Wang C, Fu Y, Ren Z (2022) Understanding the drivers of woody plant diversity in urban parks in a snow climate city of China. J Forest Res 34:1021–1032. 10.1007/s11676-022-01535-9

Mireles F, Davila JI, Pinedo JL, Reyes E, Speakman RJ, Glascock MD (2012) Assessing urban soil pollution in the cities of Zacatecas and Guadalupe, Mexico by instrumental neutron activation analysis. Microchem J 103:158–164. 10.1016/j.microc.2012.02.00

Mitrović M, Kostić O, Miletić Z, Marković M, Radulović N, Sekulić D, Jarić S, Pavlović P (2023) Bioaccumulation of potentially toxic elements in tilia tomentosa Moench trees from urban parks and potential health risks from using leaves and flowers for medicinal purposes. Forests 14: 2204. 10.3390/f14112204Su13179784

Mitrović M, Blanusa T, Pavlović M, Pavlović D, Kostić O, Perović V, Jarić S, Pavlović P (2021) Using fractionation profile of potentially toxic elements in soils to investigate their accumulation in Tilia sp. leaves in urban areas with different pollution levels. Sustainability 13:9784. 10.3390/su13179784

Molnár VÉ, Simon E, Ninsawat S, Tóthmérész B, Szabó Sz (2020) Pollution assessment based on element concentration of tree leaves and topsoil in Ayutthaya Province, Thailand. International J Environ Res Public Hlth 17:5165.

Nagajyoti PC, Lee KD, Sreekanth TVM (2010) Heavy metals, occurrence and toxicity for plants: a review. Environ Chem Lett 8:199–216. 10.1007/s10311-010-0297-8

Pietrelli L, Menegoni P, Papetti P (2022) Bioaccumulation of heavy metals by herbaceous species grown in urban and rural sites. Water Air Soil Pollut 233: 141. 10.1007/s11270-022-05577-x

Roca-Perez L, Boluda R, Rodríguez-Martín JA, Ramos-Miras JJ, Tume P, Roca N, Bech J (2023) Potentially harmful elements pollute soil and vegetation around the Atrevida mine (Tarragona, NE Spain). Environ Geochem Hlth 45: 9215–9230. 10.1007/s10653-023-01591-y

Roca-Perez L, Gil C, Cervera ML, Gonzálvez A Ramos-Miras J, Pons V, Bech J, Boluda R (2010) Selenium and heavy metals content in some Mediterranean soils. J Geochem Explor 107: 110–116, 10.1016/j.gexplo.2010.08.004

Serbula SM, Kalinovic TS, Ilic AA, Kalinovic JV, Steharnik MM (2013) Assessment of airborne heavy metal pollution using Pinus spp. and Tilia spp. Aeros Air Qual Res 13:563–573. 10.4209/aaqr.2012.06.0153.

Simon E, Baranyai E, Braun M, Cserháti C, Fábián I, Tóthmérész B (2014) Elemental concentrations in deposited dust on leaves along an urbanization gradient. SciTotal Environ 490:514–520. 10.1016/j.scitotenv.2014.05.028.

Simon E, Harangi S, Baranyai E, Braun M, Fábián I, Mizser Sz, Nagy L, Tóthmérész B (2016) Distribution of toxic elements between biotic and abiotic components of terrestrial ecosystem along an urbanization gradient: Soil. leaf litter and ground beetles. Ecol Indic 60:258–264. 10.1016/j.ecolind.2015.06.045.

Simon E, Molnár VÉ, Lajtos D, Bibi D, Tóthmérész B, Szabó S (2021) Usefulness of tree species as urban health indicators. Plants 10:2797. 10.3390/plants10122797.

Simon E, Vidic A, Braun M, Fábián I, Tóthmérész B (2013) Trace element concentrations in soils along urbanization gradients in the city of Vienna. Austria. Environ Sci Poll Res 20:917–924. 10.1007/s11356-012-1091-x.

Song M, Hu C (2018) A coupling relationship between the eco-environment carrying capacity and new-type urbanization: a case study of the Wuhan metropolitan area in China. Sustainability 10:4671. 10.3390/su10124671

Ţenche-Constantinescu AM, Madosa E, Chira D, Hernea C, Ţenche-Constantinescu RV, Lalescu D, Borlea GF (2015) Tilia spp.-Urban Trees for future. Not Bot Horti Agrobo Cluj Napoca 43:259–264. DOI:10.15835/nbha4319794

Tešić M, Stojanović N, Knežević M, Đunisijević-Bojović D, Petrović J, Pavlović P (2022) The impact of the degree of urbanization on spatial distribution, sources and levels of heavy metals pollution in urban soils—a case study of the city of Belgrade (Serbia). Sustainability 14:13126. 10.3390/su142013126

Van Gestel Cam, Ortiz MD, Borgman E, Verweij RA (2011) The bioaccumulation of Molybdenum in the earthworm Eisenia andrei: influence of soil properties and ageing. Chemosphere 82:1614–1619.

Wang S, Xiong Z, Han X, Wang L, Liang T (2024) Unveiling the spatial differentiation drivers of major soil element behavior along traffic network accessibility. Environ Poll 342: 123045. 10.1016/j.envpol.2023.123045

Wang S, Zhang H (2022) Tree composition and diversity in relation to urban park history in Hong Kong, China. Urban For Urban Green 67:127430. 10.1016/j.ufug.2021.127430

Zar IH (1996) Biostatistical Analysis. Prentice Hall, N.J.

Zeb J, Tahir H, Othman A, Habeebullah TM, Saygal A, Assaggaf HM, Ahmed OB, Sultan M, Mohiuddin S, Masood SS, Mirza AZ, Hajira B (2022) Geo-environmental approach to assess heavy metals around auto-body refinishing shops using biomonitors. Heliyon 8:e08809. 10.1016/j.heliyon.2022.e08809

